# High cholesterol increases myeloma tumour burden and promotes resistance to bortezomib

**DOI:** 10.1101/2025.01.21.633916

**Authors:** Beatriz Gámez, Danielle Whipp, Srinivasa Rao, Emma V. Morris, Young Eun Park, Zeynep Kaya, Claire M. Edwards

## Abstract

Multiple myeloma (MM) is an incurable hematologic cancer where malignant plasma cells accumulate in the bone marrow (BM), with disease progression highly dependent on cellular interactions. Obesity is a major risk factor for myeloma, however the underlying mechanisms are unclear. Here we used a murine myeloma model to show that a high cholesterol diet increases local and circulating levels of low density lipoprotein cholesterol (LDL), increasing bone marrow tumour burden in vivo. Exogenous LDL induced bortezomib-specific drug resistance in metabolically stressed myeloma cells in vitro and ex vivo. RNA-seq analysis revealed bortezomib-induced changes in the cholesterol biosynthesis/homeostasis pathway that were completely reverted with LDL pretreatment. In silico analysis of patient data supported a role for cholesterol in response to bortezomib-based therapies. Targeted proteome profiling revealed changes in expression of the adipokine resistin in the bone marrow of cholesterol-treated or myeloma-bearing mice, with elevated resistin expression observed in MGUS patients. Bone marrow adipocytes were found to be a major source of resistin, which was further increased in response to LDL. In summary, we demonstrate that high cholesterol promotes both myeloma development and bortezomib resistance, identifying resistin as a potential mediator so revealing new mechanisms underlying myeloma pathogenesis.

## INTRODUCTION

Multiple myeloma (MM) is a B cell malignancy that occurs when antibody producing plasma cells proliferate abnormally and accumulate in the bone marrow (1). MM evolves from the premalignant disorder monoclonal gammopathy of undetermined significance (MGUS), where patients show some infiltration of plasma cells into the bone marrow (≤10%) and circulating paraprotein but no end-organ damage. Although clinically asymptomatic, evidence has proven that MGUS patients suffer from bone density alterations and higher risk of fractures (2, 3). Several factors influence the progression of MGUS including the level of paraprotein and the subtype of MGUS (Ig type) (2, 4), as well as environmental factors such as obesity and aging (5, 6).

Despite recent advances in therapy and improvement in the 5-year survival rate, MM remains incurable largely due to drug resistance (7). Frequent relapses during disease ultimately exhaust any possible treatment. Among the current therapy options, proteasome inhibitors are one of the most widely used drugs in the clinic. The combination of a proteasome inhibitor together with an immunomodulatory drug is nowadays still the most effective therapy for newly diagnosed MM patients (1, 8). However, MM patients will eventually go through multiple relapses, each of them presenting shorter periods of remission as a consequence of the development of drug resistance (9).

Obesity is the second leading cause of cancer and together with excess body fat it has been linked to several types of malignancies, including MM (10, 11). Moreover, obesity and BMI (body mass index) are strongly associated with progression from MGUS to MM (5, 12, 13), with preclinical studies demonstrating increased myeloma tumour burden in response to diet-induced obesity (14, 15). Although the exact mechanism by which obesity impacts MM pathogenesis is still unclear, it is the only modifiable risk known for MM and so it provides hope for dietary and lifestyle interventions as a strategy to reduce MM progression.

Strongly associated with obesity is high cholesterol. Cells normally synthesize their own cellular cholesterol (*de novo*) through the mevalonate pathway, but they can also recruit circulating cholesterol. Circulating low-density lipoprotein-cholesterol (LDL) from diet or hepatic synthesis can interact with the LDL receptor in distant tissues. After LDL internalisation, its apolipoprotein is degraded, and free cholesterol is then available.

Several studies have shown that MM patients have a reduction in total cholesterol, LDL and high-density lipoproteins (HDL) in plasma compared to healthy individuals. Moreover, data suggests that these changes are even more evident in later stages of disease (16, 17). Increased LDL uptake by myeloma cells may explain the hypocholesterolaemia seen in MM patients. As cancer cells proliferate rapidly, they require high levels of cholesterol for plasma membrane formation and cellular functions and so can reprogram their cholesterol metabolism to satisfy these needs (18). However, despite significant efforts to elucidate the role of total cholesterol or LDL in cancer, little is known about the effect of high circulating cholesterol and LDL levels in multiple myeloma disease.

Here, we used the 5TGM1 preclinical murine model of myeloma together with a high cholesterol diet to investigate the impact of elevated cholesterol within the myeloma bone marrow microenvironment. We subsequently investigated the mechanisms by which cholesterol promotes myeloma progression and drug resistance, identifying resistin as an adipokine associated with cholesterol-tumour-bone crosstalk.

## RESULTS

### A high cholesterol diet increases circulating LDL and tumour burden in vivo

To understand the effect of cholesterol on myeloma progression, a 2% cholesterol diet was used to increase cholesterol levels in C57BL/KaLwRij mice. Mice were placed on the diet for four weeks after which time mice had an increase in circulating LDL (Fig 1A & Fig S1A). There was no significant change in body weight (Fig S1B). Signs of fatty liver disease were seen (macroscopic hepatic steatosis (data not shown) and increased liver weight (Fig S1C)). Mice were then inoculated intravenously with 5TGM1-GFP cells and distributed into two groups that had either a) cholesterol diet halted at time of tumour inoculation or b) cholesterol diet continuously given after tumour inoculation (Fig S1A). By the end of the experiment, LDL levels remained elevated only in the animals fed continuously with the cholesterol diet. LDL returned to control levels when the diet was removed at time of tumour inoculation (Fig 1B).

**Figure 1.**
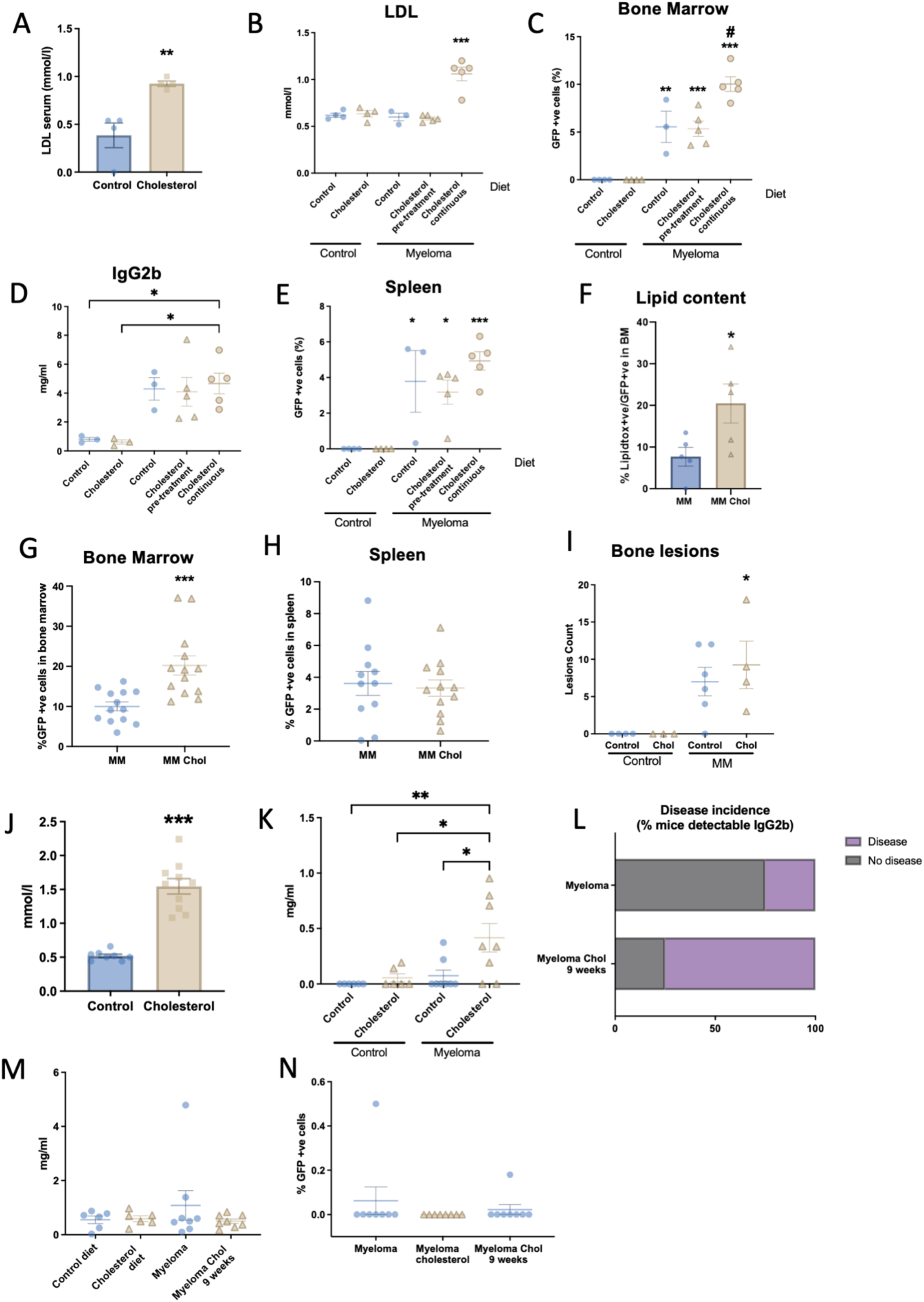
A high cholesterol diet increases LDL and bone marrow tumour burden in vivo in C57BL/6 KaLwRij mice. C57BL/6 KaLwRij mice were fed with either control or high cholesterol diet for 4 weeks and serum LDL was measured (**A**). Mice were then randomly distributed into either control or tumour-bearing mice (myeloma) and inoculated with 1.5×10^6^ 5TGM1-GFP cells. Cholesterol non-tumour mice and cholesterol pre-treatment myeloma mice had cholesterol diet halted at time of inoculation. (**B**) LDL in serum was measured at endpoint. (**C**) Proportion of 5TGM1 GFP +ve tumour cells in bone marrow. (**D**) IgG2bK levels in serum. (**E**) Proportion of GFP+ve myeloma cells in spleen. (**F**) Lipid content of GFP+ve myeloma cells from bone marrow quantified by lipidtox stain and flow cytometry. In separate experiments, cholesterol diet was given from time of inoculation and GFP+ve myeloma cells in bone marrow (**G**) and spleen tumour burden (**H**) were quantitated by flow cytometry. (**I**) Number of bone lesions counted from microCT images. C57BL/6 animals were placed on a high cholesterol diet for 5 weeks prior to tumour inoculation (1.5×10^6^ 5TGM1-GFP cells) and serum LDL measured (**J**). 4.5 weeks after tumour inoculation, IgG2bK levels (**K**) and disease incidence (% of mice with detectable IgG2b) were assessed (**L**). 6 weeks after tumour inoculation all mice were culled and IgG2bK paraprotein (**M**) and tumour burden (**N**) measured. For 2 group comparison, two-tailed Student’s t-test was performed. *p< 0.05, **p < 0.01 *and \**\*\*p < 0.001. For more than 2 groups one-way anova analysis was performed. If not otherwise indicated, *p< 0.05, **p < 0.01 and *\**\*\*p < 0.001 compared to control with no-tumour. #p < 0.05 compared to myeloma control mice. Results are presented as mean ± SEM.

All 5TGM1-GFP inoculated mice had evidence of myeloma as evidenced by the proportion of tumour cells in bone (Fig 1C) and the rise of paraprotein in plasma (Fig 1D). Only those mice continuously fed with a high cholesterol diet had a significant increase in tumour burden compared to the myeloma control group (Fig 1C). Accordingly, IgG2b levels were significantly higher than non-tumour mice only in the group continuously fed with cholesterol (Fig 1D). No significant changes in spleen tumour burden or weight were detected in any of the cholesterol diet groups suggesting a bone specific effect of LDL (Fig 1E & Fig S1D). Myeloma cells from continuously cholesterol fed animals had a higher proportion of highly lipidtox+ve cells demonstrating a higher intracellular lipid content (Fig 1F).

Thus, cholesterol diet increased myeloma only when LDL levels were high during tumour development (Fig 1B-D). To further investigate this, we studied mice where the cholesterol diet was introduced only at time of tumour inoculation (Fig S1A), observing significantly higher bone marrow tumour burden with no changes in spleen tumour burden (Fig1G-H). MicroCT analysis demonstrated that only animals fed with cholesterol diet had a significant number of bone lesions compared to control animals (Fig 1I).

Unlike C57Bl/KaLwRij mice, C57BL/6 mice are not typically permissive for 5TGM1 myeloma cell growth after tumour inoculation. However, we have previously shown that diet-induced obesity in C57BL/6 mice promotes a myeloma like condition in vivo (14) and increases circulating cholesterol (data not shown). Therefore, we tested whether a high cholesterol diet would also render C57BL/6 mice susceptible to myeloma development. C57Bl6 mice on a cholesterol diet for 5 weeks had higher circulating LDL levels with no significant changes in body weight and evidence of fatty liver (Fig 1J & Fig S1E-G). Mice on the cholesterol diet for 5 weeks before and 4 weeks after tumour inoculation showed a significant increase in the levels of IgG2b compared to tumour-bearing mice on regular diet (Fig 1K). Moreover, the incidence of disease (proportion of mice with detectable IgG2b within each group) was higher in the cholesterol fed group (Fig 1L). Due to experimental limitations, mice could only be fed with cholesterol diet for 4 weeks after tumour inoculation after which normal diet was introduced and mice were culled 2 weeks after diet removal. Bone marrow tumour burden and IgG2b levels at endpoint showed barely any detectable myeloma (Fig 1M & N).

### LDL restores lipid depletion-induced viability loss and induces bortezomib resistance in vitro and ex vivo

To further investigate the role of LDL on myeloma cells, a panel of human (JJN3, MM1S) and mouse (5TGM1-GFP) myeloma cell lines were cultured in media containing delipidated or complete FBS for 72 (JJN3, MM1S) or 96 hours (5TGM1-GFP). As expected, cells grown in media with no lipids showed a significant viability reduction (Fig 2A) and addition of LDL restored viability of all cell lines. Very-low-density lipoprotein (VLDL) had no effect on viability of JJN3 or MM1S. In 5TGM1-GFP cells, VLDL had similar effects to LDL suggesting differential sensitivities of myeloma cells to lipid depletion and cholesterol levels. In support of this, lipid depletion and metabolic stress (no FBS) caused an increase in LDL uptake, with variability in uptake between cell lines (Fig S2A-D).

**Figure 2.**
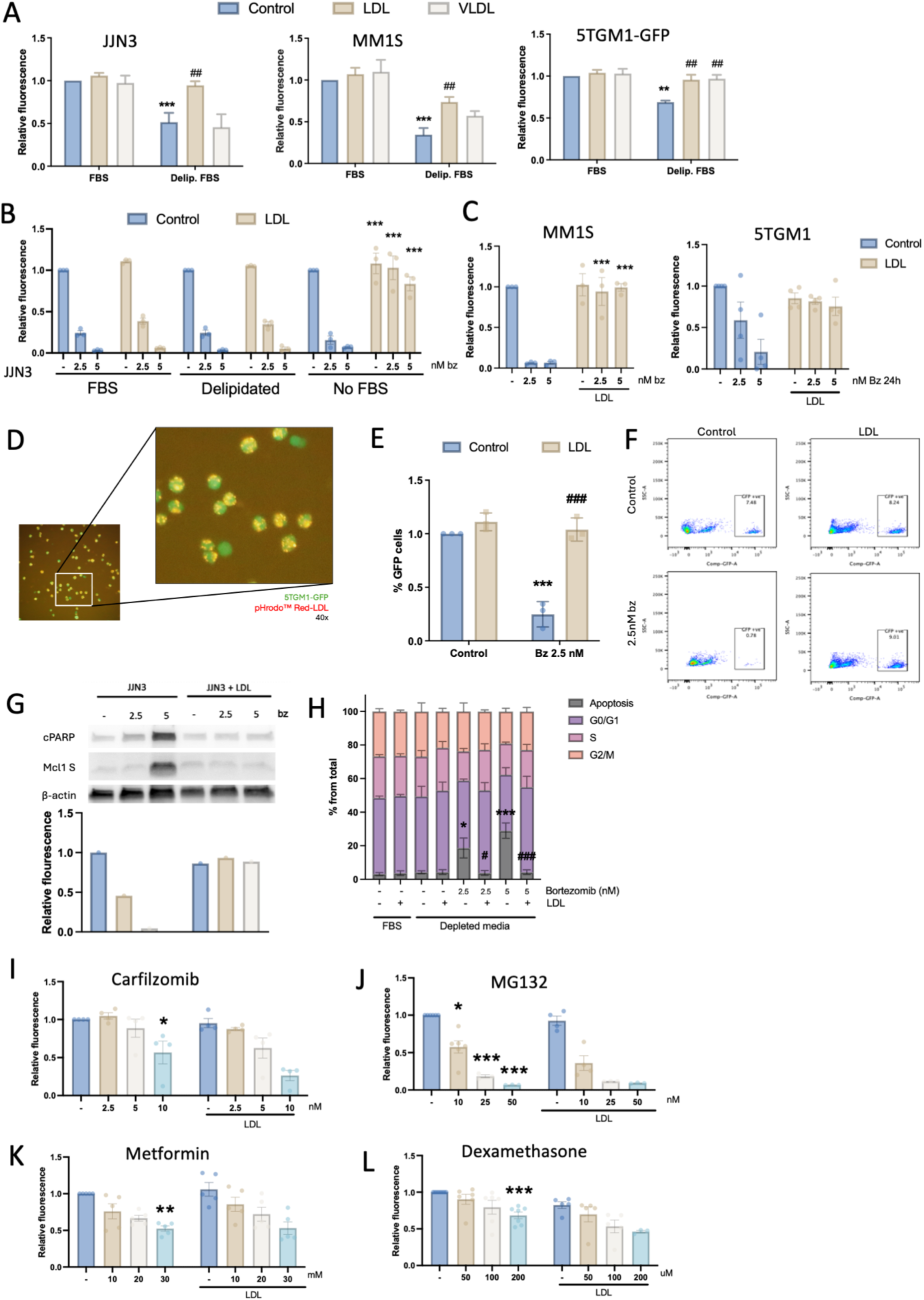
LDL can restore myeloma cell viability and induce resistance to bortezomib. JJN3, MM1s and 5TGM1-GFP cells were cultured in 10% FBS media or lipid-depleted media ± 6 µg/ml LDL or VLDL. Viability was assessed after 72 (JJN3, MM1s) or 96 (5TGM1) (**A**). JJN3 (**B**) MM1S and 5TGM1 (**C**) myeloma cells were cultured in 10% FBS containing media, delipidated media or no FBS and treated with 30 µg/ml LDL for 3 hours before 24 hour bortezomib treatment (2.5 or 5nM). Viability was quantitated by alamar blue (n = 3-7). 5TGM1-GFP were cultured in serum-free conditions and treated with pHrodo^TM^ red-LDL for 3 hours. 40x magnification (**D**). Whole bone marrow was isolated from myeloma-bearing mice and treated as for (B). Tumour cell viability was assessed by flow cytometry based on proportion of GFP+ve cells compared to control (**E**). Representative FACS plot showing proportion of GFP+ve (GFP gating) on whole bone marrow content coming from one mouse with and without LDL and bortezomib treatment ex vivo (**F**). JJN3 were cultured in serum-free conditions ± LDL for 3 hours before treatment with bortezomib (2.5 or 5nM) for 24 hours. Apoptotic markers cPARP and short Mcl-1 were measured by western blotting with concurrent viability quantitation (**G**) and cell cycle analysis using propidium iodide (**H**). JJN3 cells were treated with carfilzomib (**I**), MG132 (**J**), Metformin (**K**) or Dexamethasone (**L**) ± LDL. One-way ANOVA analysis was performed. For A **p < 0.01 and *\**\*\*p < 0.001 compared to control FBS condition. ##p < 0.01 compared to control delipidated condition. B ***p < 0.001 compared to control non-LDL condition. For C ***p < 0.001 compared to same bortezomib dose without LDL. For E ***p < 0.001 compared to control non-bortezomib treated condition, ###p < 0.001 compared to bortezomib treated no-LDL condition. For H, two-way anova was performed. *p< 0.05, ***p < 0.001 compared to not treated. #p< 0.05, ###p < 0.001 compared to same bortezomib treatment without LDL. (I-L) One-way anova analysis was performed. *p< 0.05, **p < 0.01, ***p < 0.001 compared to control (no bortezomib, no LDL). Results are presented as mean ± SEM.

Our in vivo and in vitro data demonstrate the impact of cholesterol on MM cell growth. To investigate if cholesterol could also have a role in drug resistance, we examined the effect of the proteasome inhibitor bortezomib on MM cells in different lipid culture conditions in the presence or absence of LDL. LDL addition for 3 hours before treatment with bortezomib completely blocked the effect of bortezomib to reduce viability only in metabolically stressed cells (FBS-depleted) (Fig 2B-C). We confirmed that 3h was sufficient for LDL incorporation into myeloma cells (Fig 2D). To further corroborate the protective effect of LDL, ex vivo LDL-treated bone marrow isolated from myeloma-bearing mice had the same level of protection from bortezomib as seen by the proportion of GFP+ve cells in the BM (Fig 2E-F).

### LDL protects from bortezomib-induced proteasome inhibition and apoptosis but not other drug-induced death

To further investigate the relationship between cholesterol and bortezomib resistance, apoptosis and cell cycle analysis experiments were performed. As expected, bortezomib treatment reduced viability and induced expression of apoptotic markers cleaved PARP (poly ADP-ribose polymerase) or the pro-apoptotic form of Mcl-1 (Mcl-1 S). This was reversed upon addition of LDL prior to bortezomib treatment (Fig 2G). LDL alone had no effect on the cell cycle, but restored the apoptotic fraction that was increased following bortezomib treatment. (Fig 2H). To better understand the specificity of this drug resistance effect, other drugs used in myeloma treatment were tested, including the recently approved proteasome inhibitor carfilzomib. 24-hour treatment with increasing doses of carfilzomib, MG132, metformin or dexamethasone induced at least a 50% reduction in viability (Fig 2I-L). No difference in drug response was seen when LDL was added prior to the use of any of these treatments suggesting specificity for the effect of LDL on bortezomib resistance (Fig 2I-L).

Bortezomib is a reversible proteasome inhibitor with multiple downstream effects. To further investigate how LDL induces bortezomib resistance, we studied the proteasome activity of LDL pre-treated JJN3 myeloma cells. While a reduction in proteasome activity was observed in bortezomib-treated cells, no difference was seen in myeloma cells pre-treated with LDL (Fig 3A). Accordingly, cycloheximide experiments showed that the accumulation of ubiquitinated proteins after bortezomib treatment was not present in LDL pre-treated cells (Fig 3B & C).

**Figure 3.**
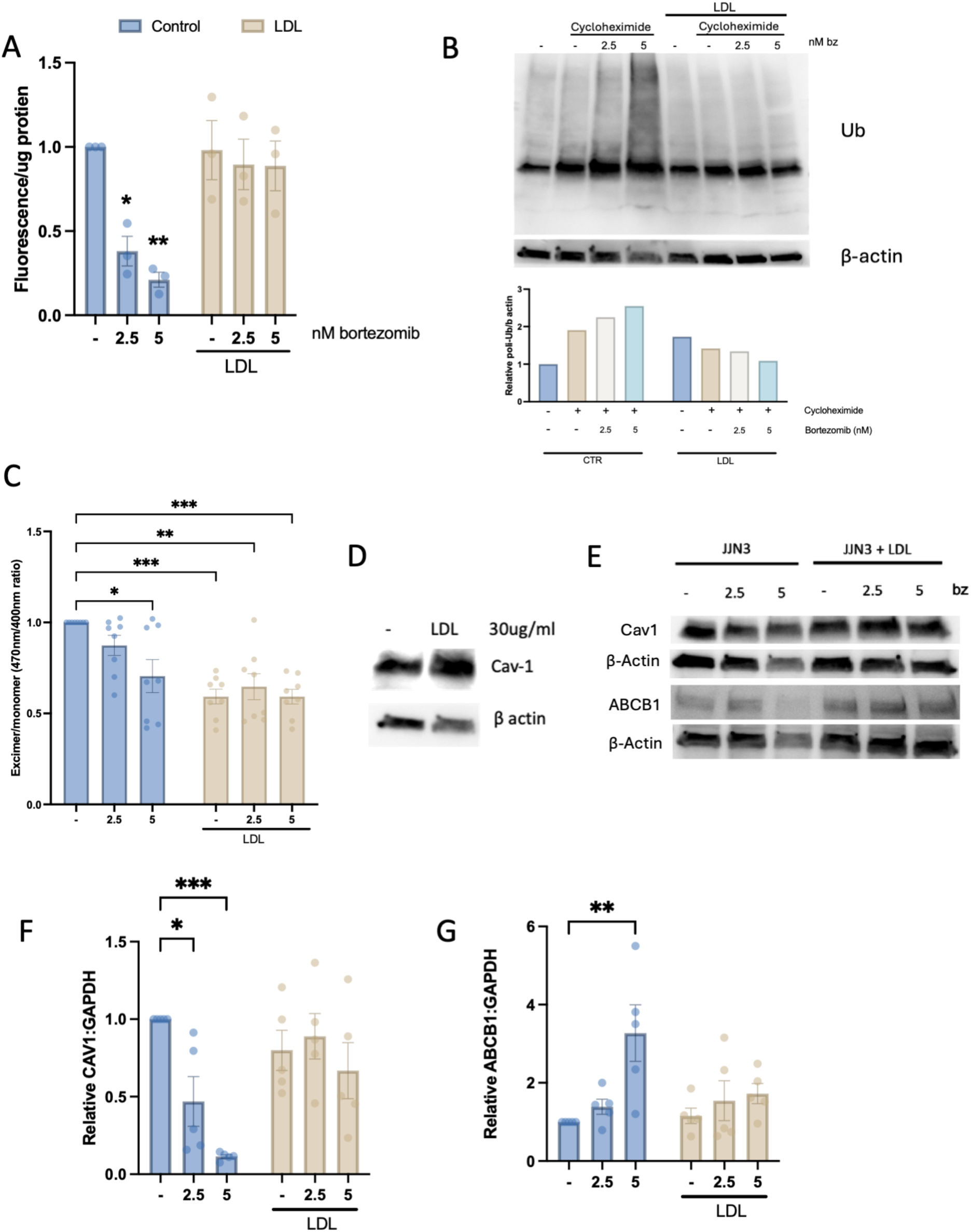
LDL treatment blocks the inhibitory activity of bortezomib and induces changes in membrane fluidity. (**A**) JJN3 myeloma cells were treated in the presence and absence of LDL and bortezomib and proteasome activity measured using a proteasome 20S activity assay kit. (**B**) Cycloheximide experiments were performed and proteasome activity assessed by expression of poli-ubiquitin. Representative western-blot and quantification (as relative ubiquitin signal/β-actin) shows ubiquitinated-protein accumulation (n=3). (**C**) Changes in membrane fluidity were evaluated. (**D**) Expression of caveolin-1 on JJN3 myeloma cells after 30μg/ml LDL treatment for 24. (**E**) Expression of caveolin-1 and MDR/ABCB1 protein in JJN3 cells treated as per drug resistance experiments. Gene expression analysis of caveolin-1 (**F**) and ABCB1 (**G**) in JJN3 cells in drug resistance experiments. One-way anova analysis was performed. *p< 0.05, **p < 0.01, ***p < 0.001 compared to control (no bortezomib, no LDL). Results are presented as mean ± SEM.

Fluctuations in cholesterol accumulation can induce changes in plasma membrane fluidity and composition (19). Bortezomib induced a dose-dependent decrease in membrane fluidity in JJN3 myeloma cells (Fig 3C). LDL pre-treatment also reduced membrane fluidity which was not further decreased by bortezomib (Fig 3C). Cellular cholesterol levels can also alter several molecular pathways associated with drug uptake, efflux and chemoresistance, including caveolin-1 and ABC transporters (20, 21). LDL increased caveolin-1 expression in myeloma cells (Fig 3D). Bortezomib-treated JJN3 cells had reduced caveolin-1 and MDR1 expression which was prevented by LDL pre-treatment (Fig 3E). Caveolin-1 gene expression was also reduced by bortezomib with reversal by LDL pre-treatment (Fig 3F). However, LDL alone did not increase cav1 (Fig 3F). MDR1 gene expression (*ABCB1*) analysis also showed differences compared to protein expression, with an increase of ABCB1 levels after bortezomib treatment that was prevented by LDL (Fig 3G).

### RNA-seq reveals that bortezomib induces changes in cholesterol homeostasis and pre-treatment with LDL is linked to ferroptosis

To further understand how LDL alters myeloma cells to overcome bortezomib treatment, RNA-sequencing (RNA-Seq) was performed on JJN-3 myeloma cells in the presence and absence of LDL and bortezomib. Differential expression gene (DEG) analysis revealed changes in expression of genes key to cholesterol synthesis (e.g. HMGCS1 or INSIG1) in response to bortezomib treatment that were prevented by LDL pre-treatment for 24h revealed (Fig 4A-B, Fig S3A). Gene set expression analysis (GSEA) revealed a significant effect of bortezomib on reactome cholesterol biosynthesis (Fig 4C) and cholesterol homeostasis and biosynthesis (Fig S3B). When GSEA hallmark gene sets were analysed in bortezomib-treated cells compared to LDL-bortezomib treated, the hallmark apoptosis gene set was enriched with bortezomib treatment (Fig S3C). Further analysis using the ShinyGO tool and top 1000 down-regulated genes demonstrated proteasome pathway enrichment in bortezomib samples (Fig 4D). Interestingly, ferroptosis was also among the most enriched pathways, demonstrated by both ShinyGO and GSEA (Fig 4D-E). Only a small number of differentially expressed genes were found after LDL treatment compared to untreated (Fig S3D-E) with ShinyGO analysis of all DEG revealing ferroptosis as one of only 2 modulated pathways (Fig S3F). GSEA using the WP Ferroptosis gene set revealed a significant effect of LDL on ferroptosis in both control and bortezomib-treated cells (Fig 4E-F). Genes related to ferroptosis include SLC7A11 and GPX4 among others (22). RNA-seq analysis showed that bortezomib induced a reduction in GPX4 and an increase in SLC7A11 (Fig 6G). SLCA711 was also elevated in LDL-treated cells (Fig 4H).

**Figure 4.**
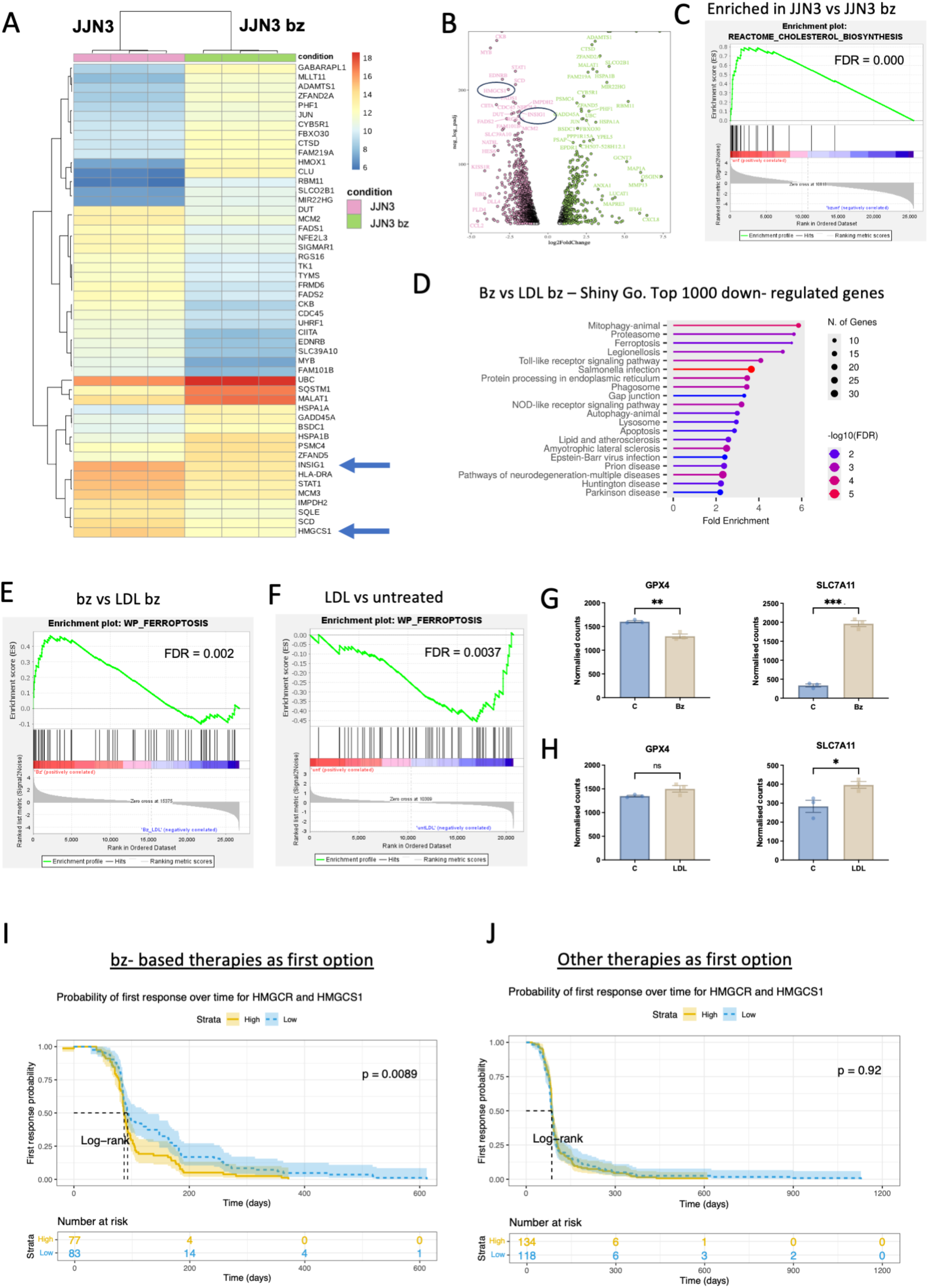
RNA sequencing reveals that bortezomib induces changes in cholesterol biosynthesis and pre-treatment with LDL is linked to ferroptosis. JJN3 myeloma cells were treated in the presence and absence of LDL and bortezomib and RNA-Seq performed. Heatmap showing differential expression gene (DEG) analysis (**A**) and volcano plot (**B**) from bortezomib vs. control cells. Pink for low and green for high expression in bortezomib samples. (**C**) Bortezomib vs control GSEA enrichment plots for cholesterol biosynthesis reactome. (**D**) ShinyGO analysis of bortezomib vs LDL + bortezomib. GSEA analysis using the WP_FERROPTOSIS gene set in bortezomib vs LDL + bortezomib (**E**) and LDL vs. control (**F**). Normalised counts of GPX4 and SLC7A11 genes in control vs. bortezomib (**G**) and in control vs. LDL (**H**). CoMMpass study data analysis showing the impact of HMGCR and HMGCS1 expression on the probability of first response over time in patients with bortezomib-based therapies as first option (**I**) and patients with other therapies (**J**). For 2 group comparison, two-tailed Student’s t-test was performed. *p < 0.05, **p < 0.01 & ***p < 0.001. Results are presented as mean ± SEM.

We next investigated whether expression of genes related to cholesterol biosynthesis was related to outcomes in bortezomib treated patients using data from the CoMMpass-MMRF database. A list of 7 mevalonate pathway-related genes (M46454 human gene set from the C2 collection-canonical pathways, GSEA) was used to study whether the expression of these genes had an impact on probability of first response in patients (Table S1). Expression of HMGCR and HMGCS1, key genes in the mevalonate pathway, appeared to have an effect in the probability of first response over time for patients that were placed under bortezomib-based therapies as first option but not for other therapies, however differences were not significant after p-value correction (Fig S3G-J). Further analysis studying the combined effect of both genes showed that they were determinant only for the response to therapy in patients under bortezomib therapy, demonstrating the importance of the cholesterol metabolism in the response to bortezomib (Fig 4I-J ).

### High cholesterol diet increases expression of resistin in bone marrow

Although we show that a high cholesterol diet increases circulating LDL, myeloma is heavily dependent on the bone marrow microenvironment, therefore we investigated the impact of dietary cholesterol on local as compared to systemic factors. VLDL/LDL cholesterol fraction in bone marrow plasma was increased in cholesterol-fed mice (Fig 5A). Using an adipokine proteome profiler we detected several differential responses, most notably the adipokine resistin which was increased with cholesterol diet or in myeloma specifically in bone marrow plasma (Fig 5B-C). Interestingly, relative to the rest of adipokines tested, resistin was lower in bone marrow plasma compared to blood (Fig S4A-B). To further corroborate the increase in resistin, we measured resistin by ELISA, demonstrating a significant increase in resistin following cholesterol diet in bone marrow plasma (Fig 5D) but not blood (Fig 5E), suggesting a specific role of cholesterol in the bone marrow plasma. Tumour-bearing mice showed significantly higher resistin in BM plasma (Fig 5F), with a positive correlation between bone marrow plasma resistin and IgG2bκ (Fig 5G).

**Figure 5.**
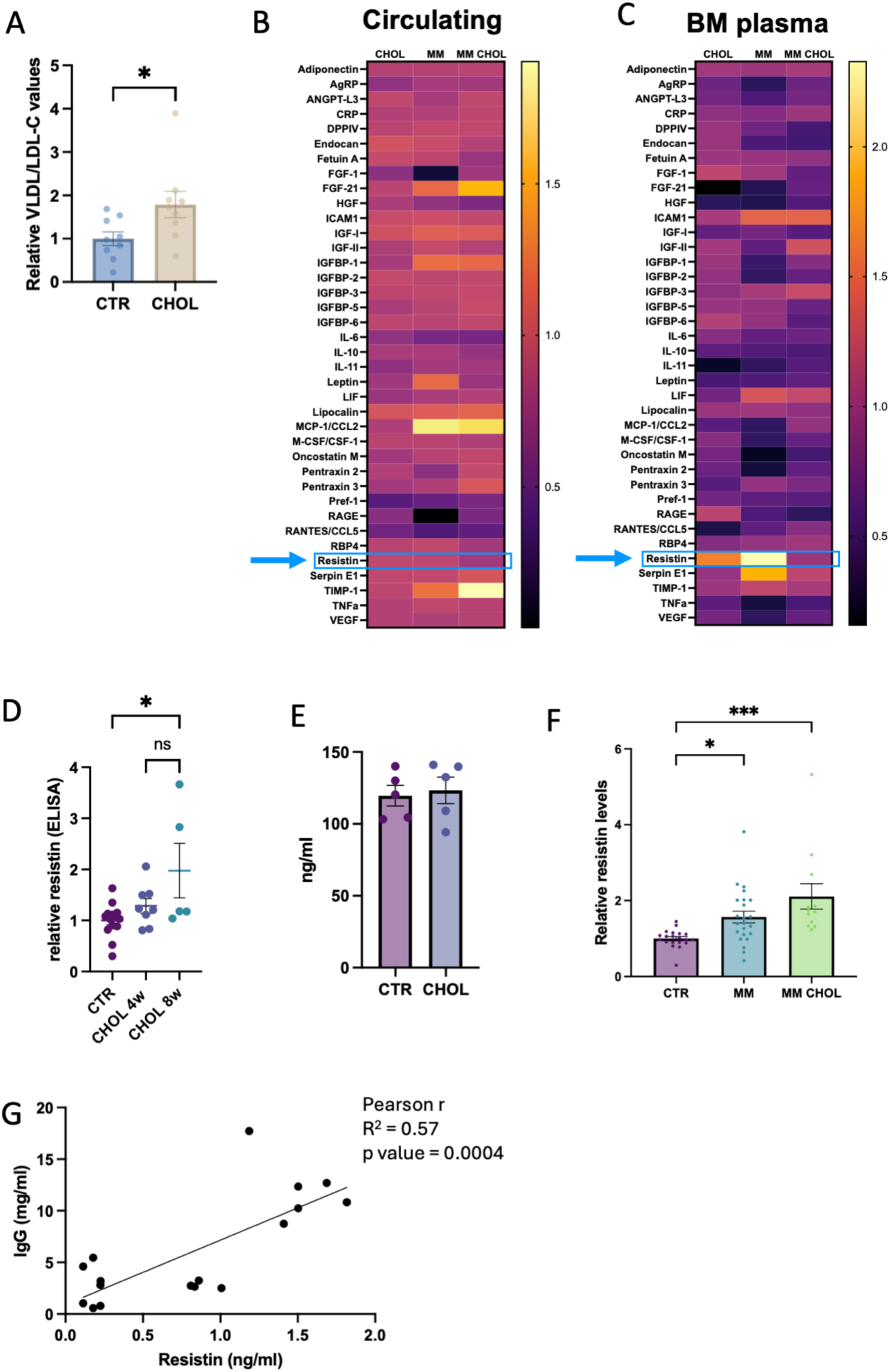
Adipokine profiles identify increased resistin in bone marrow in response to high cholesterol or myeloma. (**A**) LDL/VLDL-C ratio in the bone marrow of C57BL/6 KaLwRij mice after 4 weeks of high cholesterol diet. Adipokine analysis in blood (**B**) and bone marrow plasma (**C**). Heat maps show relative expression of adipokines in a non-tumour mouse on cholesterol diet (CHOL), myeloma-bearing mouse (MM) and myeloma-bearing mouse on cholesterol diet (MM CHOL) compared to non-tumour control. (**D**) Relative resistin expression (expressed as fold-increase compared to control) in bone marrow plasma from non-tumour mice fed with control diet (CTR), cholesterol diet for 4 weeks (CHOL 4w) or cholesterol diet for 8 weeks (CHOL 8w). (**E**) Resistin expression in blood from control or cholesterol fed animals for 8 weeks. (**F**) Relative resistin expression (expressed as fold-increase compared to control) in bone marrow plasma in control non-tumour mice (CTR), myeloma-bearing mice (MM) and myeloma-bearing mice on cholesterol diet (MM CHOL) from inoculation. (**G**) Pearson correlation of IgG2bK paraprotein and resistin levels with 17 pairs of XY coming from control and tumour-bearing mice. For 2 group comparison, two-tailed Student’s t-test was performed. For more than 2 groups one-way anova analysis was performed. If not otherwise indicated, *p< 0.05 and ***p < 0.001 compared to control with no-tumour. Results are presented as mean ± SEM.

To further study the role of resistin in myeloma pathogenesis, in silico analysis was performed (GSE47552 data (23)) with resistin levels significantly increased in CD138-positive cells from MGUS patients compared to normal plasma cells, SMM (smouldering myeloma) and myeloma (Fig 6A). In light of these results, we hypothesised whether the levels of resistin could be linked to the ability of myeloma cells to respond to bortezomib. Although a small data set, in silico analysis revealed a trend towards an increase in resistin in bortezomib-resistant patients, which was significantly increased with ex-vivo bortezomib treatment (GSE51940 (24)) (Fig 6B).

**Figure 6.**
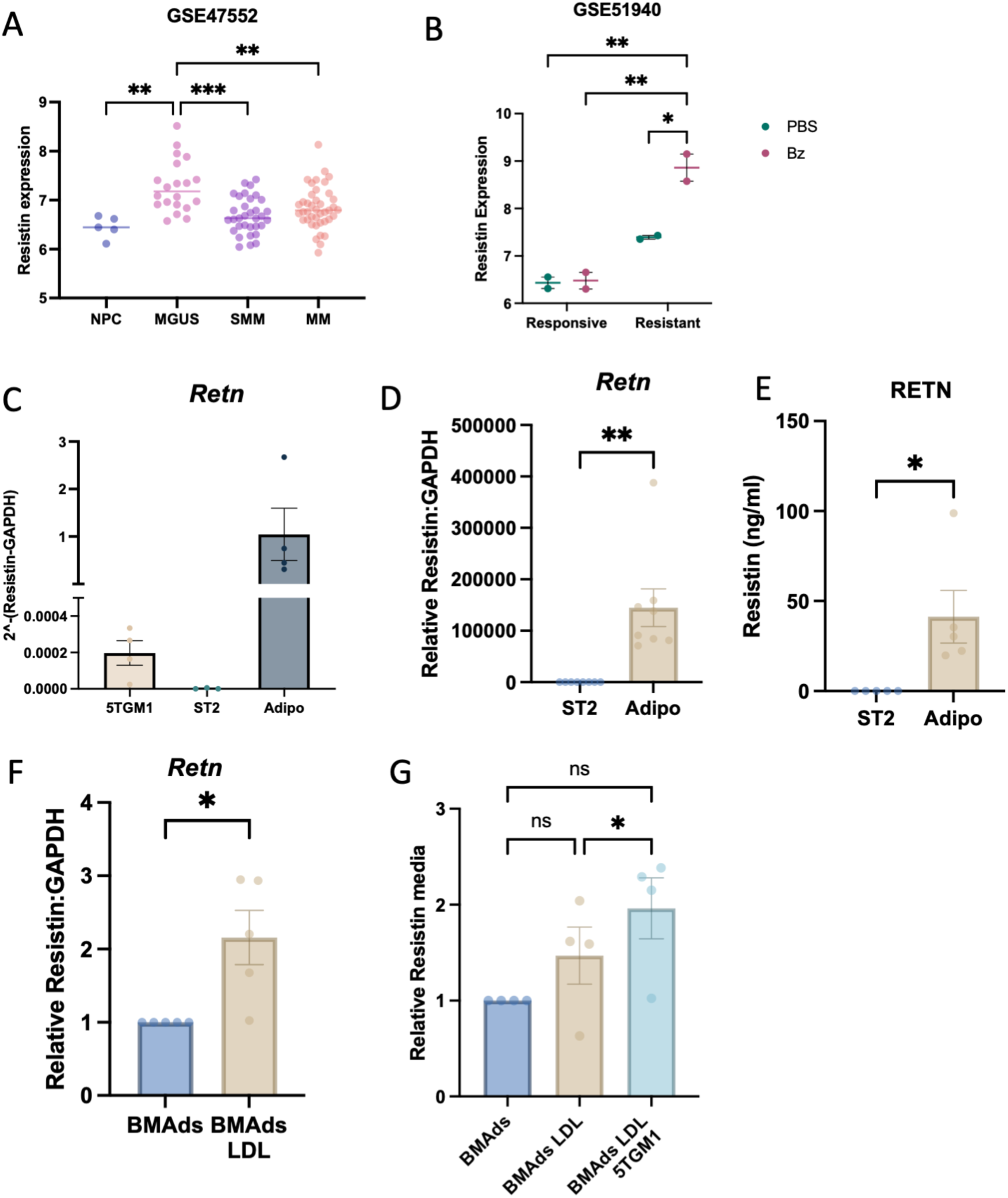
Resistin is increased in patients with MGUS and in bone marrow adipocytes following LDL treatment. (**A**) CD138+ve plasma cells from GSE47552 dataset were analysed using GEO2R and resistin expression is shown. (**B**) Whole bone marrow from bortezomib-responsive and bortezomib-resistant patients (GSE51940 dataset) was analysed using GEO2R and resistin expression is shown. (**C**) Resistin gene expression in 5TGM1, ST2 and ST2-derived BMAds (Adipo). (**D**) Relative resistin gene expression comparing undifferentiated ST2 (ST2) and ST2-derived BMAds (Adipo). (**E**) Resistin protein concentrations secreted by undifferentiated ST2 and ST2-derived BMAds (Adipo). (**F**) Relative resistin expression in BMAds compared to BMAds treated for 3 days with exogenous LDL. (**G**) Relative resistin expression in media from BMAds when treated with LDL or cocultured with 5TGM1 + LDL compared to BMAds on their own. (**H**) 5TGM1 myeloma cell line was cultured alone or cocultured with undifferentiated ST2 or BMAds and resistin gene expression was quantified. For 2 group comparison, two-tailed Student’s t-test was performed. *p< 0.05, **p < 0.01. For more than 2 groups one-way anova analysis was performed. If not otherwise indicated, *p< 0.05, **p < 0.01 & ***p < 0.001. Results are presented as mean ± SEM.

### LDL increases resistin secretion from bone marrow-derived adipocytes

Given that resistin is an adipokine, and that we have previously shown that BMAds promote myeloma development (25), we next investigated the effect of LDL on resistin expression in myeloma-bone crosstalk. BMAds (differentiated from ST2 bone marrow stromal cells) expressed high resistin, approximately 5×10^4^ times more than 5TGM1 myeloma cells, whereas undifferentiated ST2 bone marrow stromal cells did not express resistin (Fig 6C-D). Similarly, BMAds secreted resistin whereas no detectable levels were found in matched ST2 cultures (Fig 6E). LDL-treated BMAds had increased resistin gene expression (Fig 6F). We next treated BMAds with LDL either alone or in transwell coculture with 5TGM1 myeloma cells. LDL treatment of the BMAd/5TGM1 coculture significantly increased secreted resistin (Fig 6G). Taken together, these results suggest that BMAd-derived resistin may play a role in the effect of cholesterol on myeloma progression.

## DISCUSSION

Cholesterol levels have been associated with myeloma progression, with the widely used cholesterol-lowering drugs statins reported to have anti-myeloma effects in vitro and associated with progression from MGUS to myeloma (26–29). However, despite this evidence there is a limited understanding of the complex mechanisms underlying the effect of cholesterol in the bone marrow niche which is vital in order to fully exploit the prognostic and therapeutic potential of this metabolic pathway. In this study, high dietary cholesterol increased myeloma tumour burden specifically in the bone marrow. This was associated with an increase in lipid content of myeloma cells, suggesting myeloma cells uptake cholesterol to support survival. Mechanistic studies revealed that LDL-cholesterol induced resistance to the proteasome inhibitor bortezomib, with transcriptomic profiling revealing the impact of bortezomib on cholesterol metabolism and implicating ferroptosis in mediating drug resistance. Proteome profiling of the local bone marrow microenvironment identified the adipokine resistin as elevated in both high cholesterol and myeloma, and as new mediator of the tumour-bone crosstalk driving myeloma pathogenesis.

A high cholesterol diet increased myeloma tumour burden in the 5TGM1 murine model of myeloma, which relies upon inoculation of 5TGM1 myeloma cells into C57Bl/KaLwRij mice. In contrast to C57Bl/KaLwRij mice, C57BL/6 mice can recapitulate signs of multiple myeloma only under certain situations that make them prone to disease as is the case of diet-induced obesity, myeloma-like model reminiscent of MGUS (14). In the present study, when C57Bl6 mice were placed on a high-cholesterol diet, at endpoint, no features of myeloma were observed, suggesting that unlike diet-induced obesity, high cholesterol is not sufficient to alter the bone marrow microenvironment to render non-permissive mice permissive to myeloma.

Interestingly, the cholesterol-fed myeloma-bearing group presented a significant increase in paraprotein levels and in disease incidence 4 weeks after tumour inoculation compared to myeloma-bearing mice on a control diet. However, at endpoint levels of paraprotein or tumour burden were not significant in any of the groups. Due to our experimental setting at that time, diet was halted at 4 weeks. Recent studies have reported that C57BL6 mice show a stronger initial anti-myeloma immunity compared to C57BL/KaLwRij where they are able to limit the tumour development (30). In considering these results together with the C57Bl/KaLwRij model, where effects in cholesterol diet were seen only in consistently high LDL animals, we can speculate that removal of the diet stopped the pro-tumourigenic effect in C57BL6 mice and/or that immunity helped to detain a further development of the tumour. Taken together, our in vivo data suggests a pro-tumorigenic effect of cholesterol. This may come from a direct effect of cholesterol on tumour cells or in response to rapidly fluctuating changes in the bone marrow niche that favour the progression of myeloma cells.

In line with previous studies, our findings show that exogenous LDL is able to restore viability when myeloma cells are grown under lipid-depletion conditions (31). VLDL had no effect on JJN3 or MM1S but had similar effects to LDL on 5TGM1. Additional experiments confirmed that myeloma cells can efficiently uptake LDL, especially under metabolic stress (FBS depletion) with variability between cell lines. This agrees with previous results from our group where JJN3 and 5TGM1 showed a marked differential ability for lipid uptake (25).

High LDL availability prior to bortezomib treatment completely abrogated bortezomib-induced apoptosis in a range of myeloma cell lines. This protection was also seen in myeloma cells ex-vivo. Bortezomib-induced apoptosis was completely restored as seen by viability experiments, downregulation of apoptotic markers and restoration of the apoptotic fraction. RNA-seq also showed LDL completely reverting the bortezomib gene expression signature. Interestingly, this effect on drug resistance is exclusively seen under metabolic stress and not under standard culture conditions or using lipid depleted media. Although this might seem an unexpected finding, metabolic reprogramming represents one of the hallmarks of cancer cells contributing to cell heterogeneity, progression, and drug resistance. It is known that serum deprivation can prime cells to adapt to death supporting long-term survival in cancer cells (32). Interestingly, LDL was not able to protect from other drugs such as other proteasome inhibitors (carfilzomib and MG132), metformin or dexamethasone. More investigation is needed to fully understand the exclusive protection of LDL with regards of bortezomib. However, mounting evidence already probes different responses to closely related proteasome inhibitors, with bortezomib showing distinct cell toxicity and resistance to carfilzomib in vitro and in the clinic, where carfilzomib is an option to overcome bortezomib resistance in patients (33, 34).

Our in vitro data demonstrates that LDL pre-treatment of myeloma cells prevents the proteasome inhibitory function of bortezomib. Moreover, LDL induces changes in the membrane fluidity and expression of caveolin-1, a protein involved in cell membrane maintenance and drug resistance. Cholesterol is a crucial component of cell wall lipids, essential for formation of lipid rafts and caveolae (35). Previous studies have shown how cancer cells can exhibit lower plasma membrane fluidity leading to excessive signalling and promoting tumour growth and progression (36). Cholesterol can regulate membrane fluidity regulation and modulate intracellular signalling, cellular trafficking, and chemotherapy resistance (19, 37). Cholesterol has been previously described to increase the levels and membrane distribution of caveolin-1 protein (21).

Caveolin-1 has been shown to regulate redox homeostasis, cell adhesion and resistance to bortezomib in myeloma cells (38). Our results showed that caveolin1 increased after LDL treatment. When compared to myeloma cells treated with bortezomib, LDL pre-treated cells before bortezomib addition showed a significant increase in cav-1 gene expression and protein levels. Similar results were seen regarding the multidrug resistance MDR1/ACB1 transporter. Modest changes in the expression of these proteins may be due to assessing whole lysates instead of membrane extracts. Further work is needed to better understand the role of these proteins in cholesterol-induced bortezomib resistance.

To further understand the link between cholesterol pathway and bortezomib resistance in patients, in silico analysis using the MMRF CoMMpass study data was performed. The level of expression of key genes in the mevalonate pathway HMGS1 and HMGCR determined the first response probability over time only in patients with bortezomib-based therapy as a first option but not when other therapies were given therefore corroborating a role for cholesterol in the ability to respond to bortezomib. This is supported by reduced plasma cholesterol and LDL levels in patients achieving best-response to bortezomib in less than 66 days as compared with late responders (39).

RNA-seq analysis demonstrated that bortezomib treatment affects the expression of genes involved in cholesterol synthesis and that LDL pre-treatment was able to restore most of the pathways that were originally affected by bortezomib treatment. Enrichment analyses highlighted a decrease in ferroptosis as a possible pathway involved in the protective effect of LDL. Ferroptosis is a form of cell death that occurs as a consequence of metabolic dysregulation and is characterized by iron overload, reactive oxygen species (ROS) accumulation and lipid peroxidation (40). Changes in cholesterol homeostasis have been linked to resistance to ferroptosis with studies showing that statins can induce ferroptosis (19, 41). Moreover, the ferroptosis pathway has been proposed as an anticancer strategy to overcome drug resistance (42, 43). SLC7A11 is known to tightly regulate ferroptosis, with levels associated with cancer cells’ response to oxidative stress (44). Whereas high levels of overexpression can lead to increased death under oxidative stress, moderate levels are beneficial for cancer cells (44). In the current study SLC7A11 was increased 4-fold after bortezomib treatment but a much lower induction was seen in LDL treated cells compared to untreated. Taken together, these results suggest that metabolically stressed myeloma cells might benefit from LDL treatment by moderately increasing the levels of SLC7A11, thus being beneficial for the oxidative effects of bortezomib treatment, protecting the cells from ferroptosis. In support of our findings, Zhao et al. have also shown a protective effect of LDL-cholesterol in ferroptosis in other cancer cells, mediated through reduced membrane fluidity (19).

The high cholesterol diet elevated myeloma tumour burden in the bone marrow but not the spleen in vivo, suggesting an important role of the bone microenvironment. While we observed differences in both systemic and local factors, the adipokine resistin was increased specifically in the bone marrow niche in response to high cholesterol. Differences in circulating and bone marrow resistin could arise from differences in the lipid metabolism of BMAds and subcutaneous adipocytes, with BMAds known to display a cholesterol-orientated metabolism (45). Resistin is predominantly expressed by adipocytes, and other than its link with insulin resistance (hence the name resistin) it has been described to have a role in inflammation and cancer development (46, 47). Moreover, resistin has been linked to MM for its ability to induce multidrug tolerance (including bortezomib) and to accelerate drug expulsion by upregulating ABC transporters (48). Here we show how bone marrow derived adipocytes (BMAds) differentiated from ST2 express a vast amount of resistin. We further show that BMAds exposure to LDL increases resistin secretion which intensifies in the presence of myeloma cells. Although myeloma cells express almost no resistin, coculture of these cells with ST2 increased their resistin levels. Taken together, these results suggest that resistin levels in bone marrow plasma could be influenced by multiple factors. Firstly, the presence of high levels of LDL locally, which may change the levels of resistin secreted by the BM adipocytes. Secondly, myeloma cells colonizing the bone marrow and interacting with stromal and BMAds could contribute to the resistin pool. In silico analysis revealed that CD138+ve cells from MGUS have significantly higher resistin compared to normal plasma cells but not in MM supporting the idea of BM/tumour fluctuations. This could reflect distinct resistin expression levels when different levels of myeloma bone marrow infiltration occur. We have previously shown that BMAd number increase with low bone marrow infiltration (<10%) but subsequently decreases with high bone tumour burden (>30% infiltration) where BMAds localise to the tumour-bone interface (25). Thus, we hypothesize that resistin levels may be dependent upon extent of myeloma infiltration and proximity to stromal/BMAds. In silico analysis (GSE51940) found no significant increase in resistin gene expression in bortezomib-resistant myeloma patients, but interestingly ex-vivo bortezomib treatment of these cells increased resistin expression. Unlike the GSE47552 dataset, these samples were whole bone marrow content so reflect the contribution of all bone marrow cells in resistin expression. Further investigations are needed to fully understand the myeloma-bone cellular crosstalk impact on resistin and its role in myeloma development and drug resistance.

In summary, our findings reveal the impact of high cholesterol on myeloma progression and bortezomib resistance and provide insight into the cellular mechanisms that underly this, identifying resistin as a potential mediator within the tumour-bone microenvironment. Obesity, of which high cholesterol is a key component, is a major risk factor for myeloma. While cholesterol can be easily monitored in patients, an improved understanding of the mechanism of action of cholesterol in myeloma is required to fully exploit such data. In particular, it is intriguing to speculate whether cholesterol-lowering drugs would be beneficial for those patients receiving bortezomib. Furthermore, our studies provide important mechanistic insight to facilitate optimal pharmacological or dietary intervention strategies.

## METHODS

### Animal experiments

All procedures were conducted in accordance with the Animals Scientific Procedures Act of 1986 (UK) and approved by the University of Oxford Animal Welfare and Ethical Review Body (Home Office Project Licenses PCCCC8952 and PP9500304). 4-6 week old C57BL/6 or C57BL/KaLwRij (Inotiv) mice were used. Within each study, mice were age, sex and weight matched between treatments. Mice were housed under standard housing, husbandry, and diet conditions. Mice were randomly allocated to experimental groups. For cholesterol diet experiments 2% cholesterol diet was used (TD.0784, Inotiv) with matched control diet (T.2916MI.12, Inotiv). For cholesterol pre-treatment, mice were fed with cholesterol diet for 4 weeks prior to tumour inoculation when diet was halted. Cholesterol continuous group was fed with the diet for 4 weeks before tumour inoculation and continued after until sacrifice. As previously published (49), mice were inoculated via tail injection with 1.5 × 10^6^ 5TGM1-GFP cells and sacrificed at day 25 post-inoculation. Bone marrow was flushed, and spleen crushed. Cells were then filtered (70 µm filter) and analysed for percentage GFP fluorescence by flow cytometry. Serum <u>IgG2bK</u> concentration was quantified by <u>ELISA</u> (Mouse <u>IgG2b</u> ELISa quantification Set, Bethyl Laboratories or 88-50430-86, Invitrogen). Resistin concentrations were quantified in bone marrow and serum using mouse resistin ELISA kit (EMRETN, Invitrogen).

Circulating LDL levels were quantitated at the Clinical Pathology Laboratory at the Mary Lyon Centre (MRC Harwell) and VLDL/LDL quantifications from BM with the Cholesterol Assay Kit ab65390 (abcam).

For ex vivo drug resistance experiments, whole bone marrow content was flushed and treated with red blood cell lysis buffer, then filtered (70 µm) and analysed by flow cytometry to get time 0 percentage of GFP fluorescent cells (tumour cells). Then, cells were seeded into 96 well plates, treated with LDL for 3 hours and bortezomib (2.5nM) was added for 24 hours more. Then cells were analysed for flow cytometry again to check the proportion of myeloma cells (% GFP) after treatment in both conditions.

For lipidtox experiments, lipidtox was added to whole bone marrow content (2:1000) and left 30 minutes at 37°C before flow cytometry analysis. Gating for GFP+ve/Lipidtox high +ve gating was performed. Data was analysed using FlowJo software.

### Cell culture

5TGM1 murine myeloma cells were a kind gift from Prof. Gregory Mundy, University of Texas Health Science centre at San Antonio . MM.1S (ATCC CRL-2974) cell line was a gift of Prof. Udo Oppermann, University of Oxford. JJN-3 (DSMZ, ACC 541) and MM1-S (ATCC CRL-2974) cell lines were a gift of Prof. Udo Oppermann. 2T3 mouse preosteoblasts were a kind gift from Dr. Steve Harris, University of Texas Health Science centre at San Antonio. ST2 BMSCs were purchased from Riken cell Bank. All cell lines tested negative for mycoplasma. 5TGM1-GFP, JJN3 and MM.1S-GFP cells were cultured in RPMI-1640 medium supplemented with 10% foetal bovine serum (FBS), 2mM L-glutamine (G7513, Sigma), 100 U/ml Penicillin/Streptomycin (P/S, 15,140-122, Gibco), 0.1 mM MEM non-essential amino acids (M7145, Sigma) and 1mM sodium pyruvate (S8636, Sigma). ST2 cell line was cultured in DMEM with 10% foetal bovine serum (FBS), 1% L-glutamine and P/S. All cells were incubated at 37°C with 5% CO_2_. Delipidated FBS was purchased from PAN Biotech UK.

Cell viability was assessed using Alamar Blue (0.1 mg/ml). Quantification was performed on a BMG labtech FLUOstar plate reader. When indicated, an Incucyte® was used for live cell imaging. For myeloma experiments, proportion of GFP-positive myeloma cells were quantified using Incucyte® analysis software following the recommended protocol. For red LDL-uptake, red fluorescent positive confluency was compared to bright field confluency.

### In vitro bortezomib experiments

For drug resistance experiments, MM cells were seeded in the appropriate media (10% FBS, delipidated or FBS-free media) and 30µg/ml LDL (LP2-2MG) was added. After 3 hour pre-treatment, bortezomib was added at the indicated concentrations (2.5nM and 5nM were typically used). This setting was used for the assessment of apoptotic markers, cell cycle analysis, viability, proteasome inhibitor experiments, membrane fluidity assays and RNA-seq.

For LDL uptake experiments, Low Density Lipoprotein from Human Plasma, pHrodo™ Red-LDL Conjugates were used (L34355, Thermo Fisher).

For membrane fluidity assay, Membrane Fluidity Assay Kit ab189819 was used. Methyl-β-cyclodextrin (MCD) was used one hour before end of experiment as a control (C4555-1G, Merck-Sigma).

For cell cycle experiments, MM cells were spun down, washed with PBS and fixed with cold 70% ethanol then incubated at least 30 minutes at 4°C. Then cells were washed, spun at 850g and pellet diluted in 50µl RNase treated (100ug/ml) for 15-30 minutes. 200 µl PI was added (from 50µg/ml solution) and left for 20 minutes in the dark before analysis by flow cytometry.

Proteasome activity assays were performed using the Proteasome 20S Activity Assay Kit (MAK172, Merck). For cycloheximide experiments, cycloheximide (C4859, Sigma-Aldrich) was used at 10μg/ml 4-5 hours before end point.

### RT-qPCR and RNA sequencing

<u>RNA</u> isolation and DNase treatment were carried out using Direct-zol RNA Miniprep kit (R2052, Zymo Research) following manufacturer’s protocol. For qPCR experiments, RNA was then reverse transcribed using iScript Reverse Transcription Supermix (1708841, Bio-Rad). Mouse TaqMan probes for gapdh (Mm99999915_g1) and resistin (Mm00445641_m1) and human probes for gapdh (Hs02786624_g1), resistin (Hs00220767_m1), caveolin-1 (Hs00971716_m1) and ABCB1 (Hs00184500_m1) were used (Thermo Fisher Scientific).

RNA sequencing was performed as described previously by Rao SR et al.(50). RNA sequencing libraries were prepared with the NEBNext Ultra II RNA Library Prep Kit for Illumina (E7770S, New England Biolabs) and three replicates from each condition were sequenced using the Illumina NextSeq 500 as 75bp paired-end reads. Reads were aligned to the reference human genome (hg38) using the STAR aligner. Differential expression analysis carried out using DESeq2 (v1.38.3).

### Bioinformatic analysis

GSE47552 and GSE51940 were accessed through Gene Expression Omnibus – NCBI and analysed using GEO2R on the same platform.

Normalised counts from our RNA-seq data comparing JJN3 treated with bortezomib with or without LDL pre-treatment were used to run GSEA to identify significantly enriched pathways using MSigDB collections. Gene sets were considered significant at FDR < 25% and a gene set size filter of minimum 15 and maximum 500.

For ShinyGO analysis v0.81 and v0.741 were used. Gene lists of significantly differentially expressed genes (p adjusted value < 0.05) were used. Only down-regulated genes were used when specified. KEGG gene sets with an FDR cutoff of 0.05 were used for enrichment analysis.

For the CoMMpass – MRFF analysis was conducted using R (R Core Team, 2020), RStudio (Rstudio Team, 2020) and *survival* and *survminer* R packages for survival analysis. The IA17 database from CoMMpass study was used. Unstranded files derived from CD138+ cells were used. Normalised counts per million (cpm) were then filtered for low expressed genes (cpm > 0.15 in at least 1/3 of the dataset). Gene expression analysis was performed on the CD138+ve population from patients at baseline. Patient data was split into low or high expression compared to median expression and then Kaplan-Mayer curves and log-rank tests were performed. Bonferroni correction was performed for multiple comparisons. These data were generated as part of the Multiple Myeloma Research Foundation Personalized Medicine Initiatives (https://research.themmrf.org and www.themmrf.org)

### Immunoblot

Cell lysates were resolved and transferred to PVDF membranes (4561094 & 1704156, Bio-Rad). Antibodies used: β-actin at 1:5000 (A5316, Sigma), cPARP at 1:1000 (9544, Cell signaling), Mcl-1 at 1:1000 (5453, Cell Signaling), caveolin-1 at 1:1000 (3267, Cell Signaling), MDR1/ABCB1 at 1:1000 (12683, Cell Signaling) and ubiquitin at 1:1000 (58395, Cell signaling). Binding was detected with horse peroxidase-conjugated antibodies (7074 & 7076, Cell signalling, 1:5000) and images obtained with UVITEC Finealliance.

For adipokine array, Proteome Profiler Mouse Adipokine Array Kit was used following manufacturer’s instructions (ARY013, biotechne-R&D systems).

### Microcomputed tomography

Right tibiae were formalin-fixed and stored at 4 °C until needed. Bones were then mounted vertically in PBS and placed in the micro-CT scanner (Skyscan 1172 X-ray Microtomograph; Bruker MicroCT, Kontich, Belgium) performed at 37 kV/228 μA, with an isometric resolution of 9.94 μm/ pixel using a 0.5 mm aluminium filter. Reconstruction of the original scan data was performed using NRecon and analysed using SkyScan CT analyser software (Bruker). The same threshold setting for bone tissue was used for all samples. On the 3D reconstructed image, osteolytic lesions on the curved medial tibial surface were counted as bone lesions.

### Statistical analysis

Statistical analysis was performed using unpaired, two-tailed Student’s *t*-test for comparisons between two groups and one-way ANOVA for comparison of more than two groups. For all experiments differences were considered significant at *p< 0.05, **p < 0.01 *and \**\*\*p < 0.001. Results are presented as mean ± SEM.

## Acknowledgements

This research was funded by Blood Cancer UK, Rosetrees and the National Institute for Health Research (NIHR) Oxford Biomedical Research Centre (BRC). The views expressed are those of the author(s) and not necessarily those of the NHS, the NIHR or the Department of Health. We are grateful to the Multiple Myeloma Research Foundation Personalized Medicine Initiatives (https://research.themmrf.org and www.themmrf.org).

