## Supplementary information for "High cholesterol increases myeloma tumour burden and promotes resistance to bortezomib"

### SUPPLEMENTARY FIGURES

**Supplementary figure 1. Cholesterol diet does not increase body weight or spleen weight but induces signs of fatty liver.**

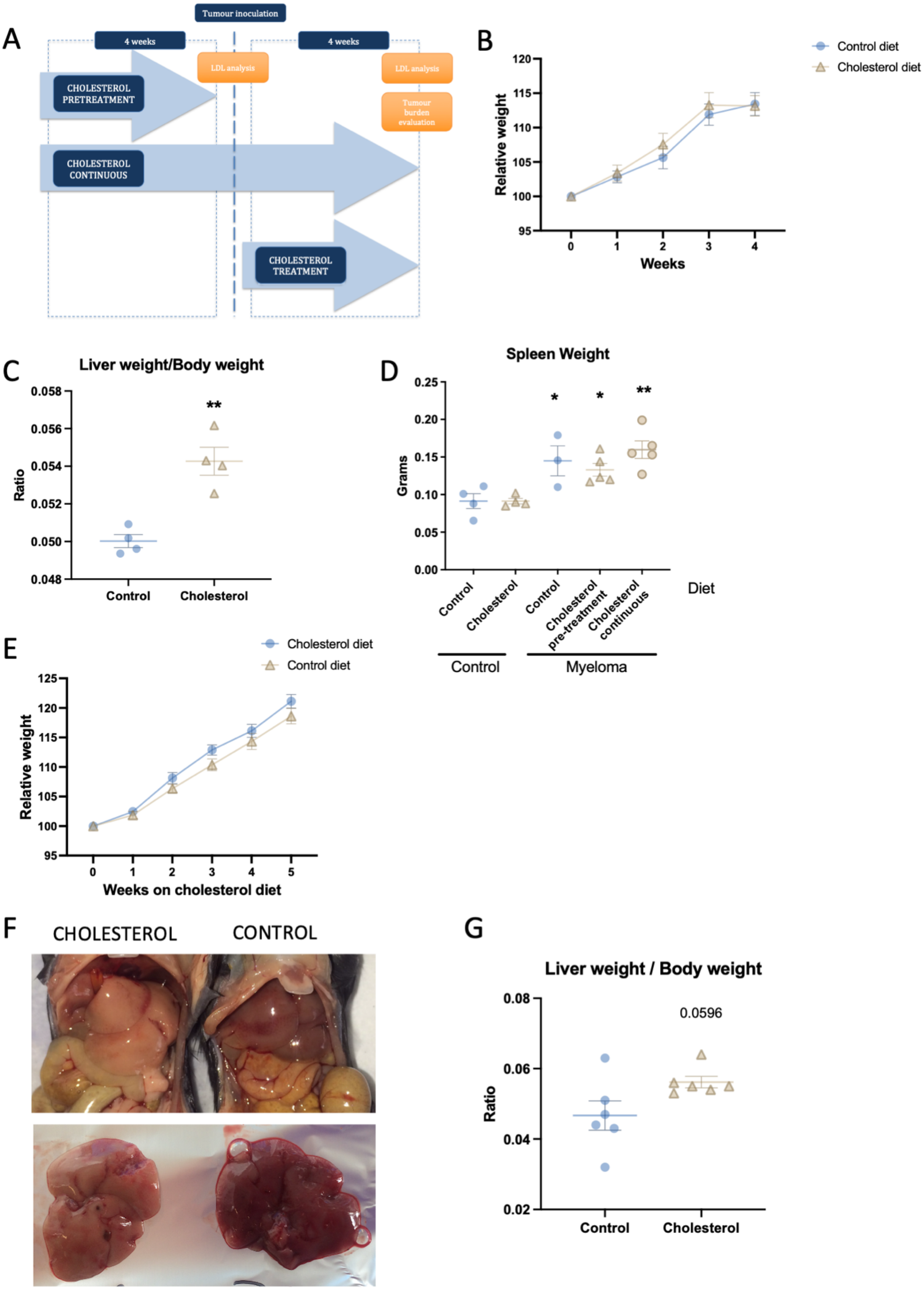

**Supplementary figure 1. Cholesterol diet does not increase body weight or spleen weight but induces signs of fatty liver.**

(**A**) Schematic figure showing the experimental setting for in vivo experiments. Mice were fed with either control or high cholesterol diet for 4 weeks and serum LDL was measured. Then animals were randomly distributed into either control or tumour-bearing mice (myeloma) and inoculated with 5TGM1-GFP cells. Cholesterol pre-treatment myeloma mice had cholesterol diet halted at time of inoculation whereas cholesterol continuous group were fed with cholesterol diet until sacrifice, where tumour burden was assessed. In separate experiments, cholesterol diet was introduced at time of tumour inoculation (cholesterol treatment). (**B**) C57BL/6 KaLwRij body weight after 4 weeks of high cholesterol diet. (**C**) Liver weight/body weight ratio in control and cholesterol-fed animals for 4 weeks. (**D**) Spleen weights where the diet was given pre-inoculation (cholesterol pre-treated) or continuously before and after tumour inoculation (cholesterol continuous). (**E**) C57BL/6 mice were fed with cholesterol diet for 5 weeks and body weight was measured. (**F**) Liver images from two representative animals under control or cholesterol diet for 5 weeks. (**G**) Liver weight/body weight ratio in control and cholesterol fed C57BL/6 for 5 weeks. For 2 group comparison, two-tailed Student's t-test was performed. $^{**}p < 0.01$. For more than 2 groups one-way anova analysis was performed. If not otherwise indicated, $^{*}p < 0.05$, $^{**}p < 0.01$ compared to control with no-tumour. Results are presented as mean ± SEM.

Supplementary figure 2. Myeloma cells differ in their ability to incorporate LDL.

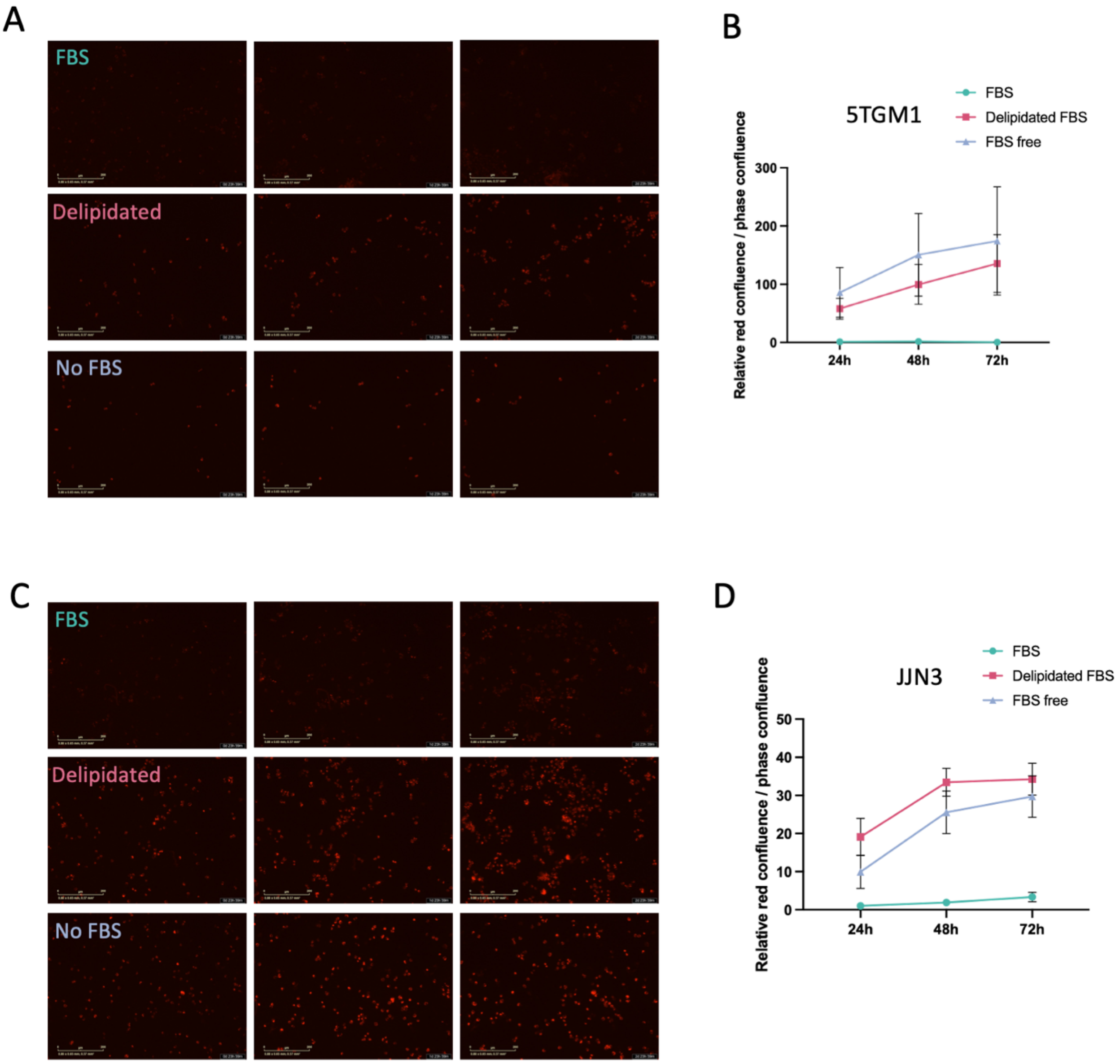


**Supplementary figure 2. Myeloma cells differ in their ability to incorporate LDL.**

5TGM1-GFP (**A-B**) and JJN3 (**C-D**) were seeded using 10% FBS media, delipidated FBS media or serum free media together with Red-LDL for 24 hours. Images were taken using Incucyte®Live-Cell analysis system (**A-B**). Quantification was performed using confluency of red signal over confluency of bright images (n=3) (**B-D**).

**Supplementary figure 3: LDL restores the gene expression signature of bortezomib treated myeloma cells.**

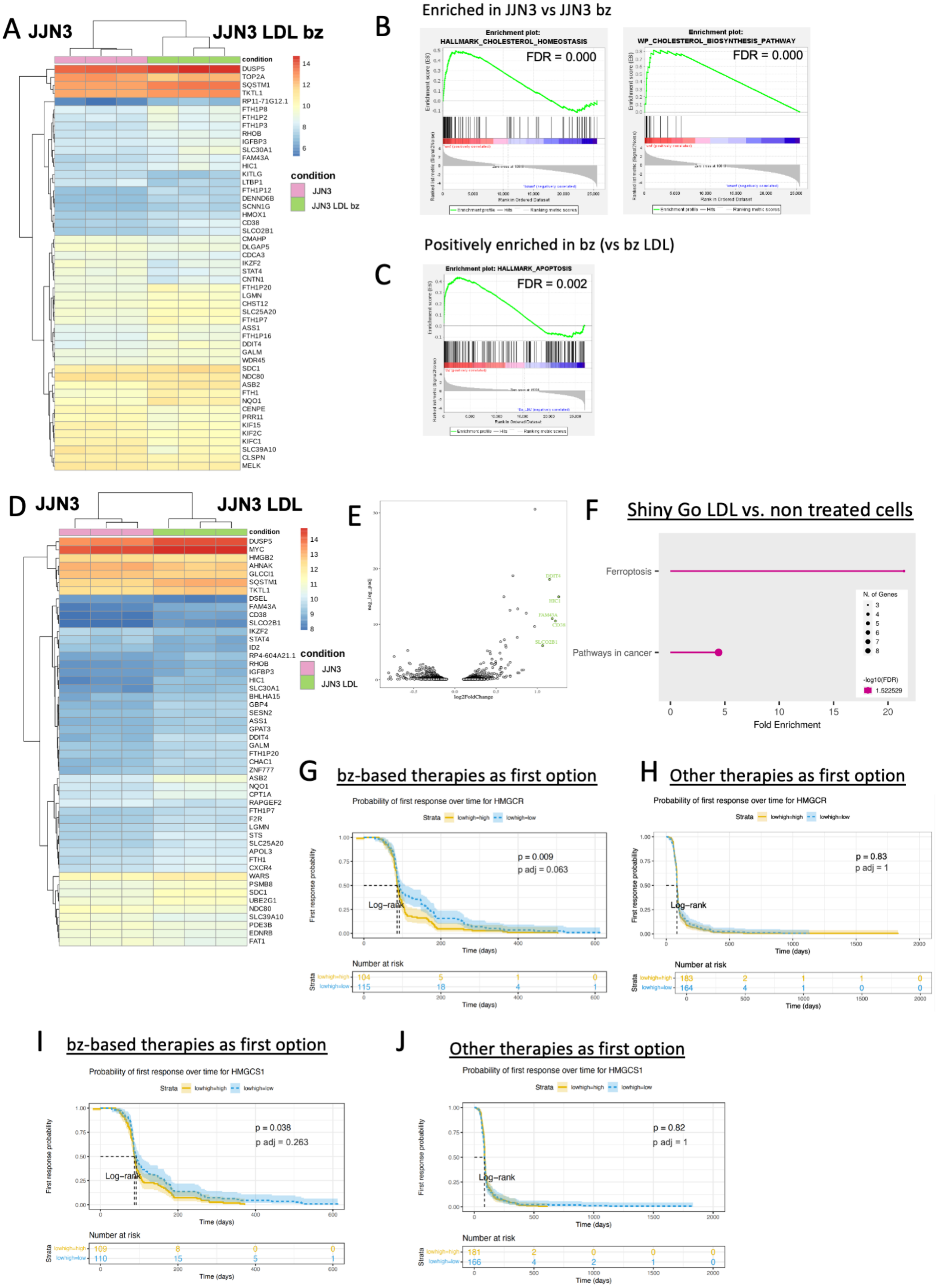

**Supplementary Figure 3. LDL restores the gene expression signature of bortezomib treated myeloma cells.**

JJN3 myeloma cells were treated in the presence and absence of LDL and bortezomib and RNA-Seq performed. (**A**) Heatmap showing DEG analysis of LDL vs. LDL + bortezomib. (**B**) Control vs. bortezomib GSEA enrichment plot of hallmark cholesterol homeostasis and WP cholesterol biosynthesis pathway gene sets. (**C**) Bortezomib vs. bortezomib + LDL GSEA enrichment plot of HALLMARK_APOPTOSIS gene set. Heatmap (**D**) and volcano plot (**E**) showing DEG analysis of LDL vs control. (**F**) ShinyGO analysis of differentially expressed genes in control vs. LDL Analysis from the MRFF-CoMMpass study data showing probability of first response over time for HMGCR (**G-H**) and HMGCS1 (**I-J**) for patients with bortezomib-based therapies as first option (**G & I**) and in patients with other therapies as first option (**H & J**). Bonferroni correction was used for adjusted p-values.

**Supplementary figure 4: Relative expression of adipokines in blood and bone marrow plasma of non-tumour mice.**

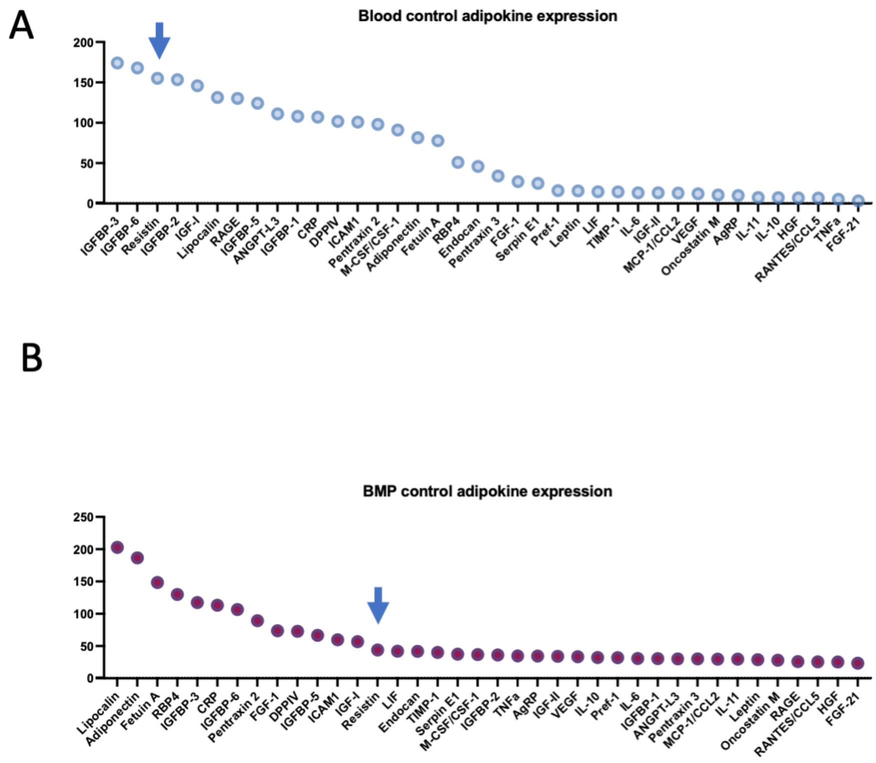


**Supplementary Figure 4. Relative expression of adipokines in blood and bone marrow plasma of non-tumour mice.**

Relative adipokine expression in blood (**A**) and bone marrow plasma (**B**) in non-tumour C57Bl/KaLWRIj mice.

### SUPPLEMENTARY TABLES

**Supplementary table 1. Mevalonate pathway genes**

| Gene symbol | Gene description |
| --- | --- |
| PMVK | phosphomevalonate kinase |
| FDPS | farnesyl diphosphate synthase |
| HMGCR | 3-hydroxy-3-methylglutaryl-CoA reductase |
| HMGCS1 | 3-hydroxy-3-methyglutaryl-CoA synthase 1 |
| ACAT2 | acetyl-Coa acetyltransferase 2 |
| MVD | mevalonate diphosphate decarboxylase |
| MVK | mevalonate kinase |

**Supplementary Table 1. Mevalonate pathway genes.**

List of genes used for analysis of CoMMpass study data. M46454 (mevalonate pathway) human gene set from the C2 collection-canonical pathways (GSEA).
